# Rapid adaptation follows experimental assisted gene flow in subset of annual monkeyflower populations

**DOI:** 10.64898/2025.12.02.689336

**Authors:** Donna M. Hinrichs, Courtney M. Patterson, Andrea Turcu, Stacy D. Holt, Simon G. Innes, Nicholas J. Kooyers

## Abstract

Assisted gene flow, the human-facilitated movement of species within its range, has the potential to increase genetic diversity and facilitate adaptation for climate-threatened populations. However, concerns regarding outbreeding depression and gene swamping have limited the application of assisted gene flow, and experimental tests have been largely confined to simulations and laboratories. We conducted a landscape-scale manipulation, planting seeds and seedlings from historically hotter and drier source populations into climate-threatened populations of the common yellow monkeyflower (*Mimulus guttatus*) and tracked change in genomes, phenotypes, and fitness. Within three generations, source alleles introgressed into half of the target populations. Assisted gene flow increased fitness in one experimental block of populations and slightly decreased fitness in a second block. Fitness increases were associated with flowering earlier and producing fewer trichomes. Greater fitness changes occurred in populations where seeds rather than seedlings were introduced. Both the amount of introgression and fitness changes were heterogeneous between years, suggesting that temporal fluctuations in climate likely impact initial introgression and success of assisted gene flow. Initial introgression from source populations was low across the genome, limiting concerns of gene swamping. Our results are generally consistent with theoretical models, and provide cautious optimism for an often-maligned conservation strategy.

## Introduction

Increasing patterns of maladaptation and declining population sizes indicate that there is a substantial threat of climate change to many organisms [1–5]. The potential risk of species extinction [4,6–9] and loss of key ecosystem services [10–14] suggest that conservation action is necessary with stipulation that actions will not further endanger species. An increasingly popular intervention is assisted gene flow (AGF) - the facilitated movement of individuals within a species range to increase the overall genetic diversity of a target population or facilitate adaptation of a target population to changing conditions [15]. Simulation and experimental evolution studies suggest that AGF could be highly successful at increasing the long-term fitness of populations [16,17], with the caveat that AGF is largely untested in natural populations [18]. Based on these potential benefits, assisted gene flow has been conducted in several plant and animal populations that are at risk of extirpation [18–21].

Although the potential of AGF is well-recognized, it is still highly controversial within the conservation community due to both perceived risks and lack of experimental data within natural populations. Two common perceived risks are outbreeding depression and gene swamping [22–24]. Introducing novel alleles into a population via outcrossing and subsequent recombination could add potentially maladapted alleles into a locally adapted population or break-up co-adaptive gene complexes [25–32]. Adding non-local alleles that are much more adaptive than local alleles could cause the loss of local genetic diversity [33,34]. Theoretical models of evolutionary rescue suggest that some maladaptation is likely to occur following introduction of new variation [35–38], but evidence of outbreeding depression is mixed with most research emphasizing that the effects of outbreeding are temporary [30,39,40]. Instead, simulation studies suggest that the benefits of AGF can be rapid (<10 generations) when fitness traits are controlled by few large-effect loci [41]. Landscape-scale tests of AGF in natural populations are necessary to evaluate potential benefits and risks.

Because AGF has historically been reserved for highly threatened species, there has been limited opportunity to dissect the consequences of AGF through controlled experiments (but see [18,42,43]. Several important hypotheses need to be evaluated. First, the impact of AGF is predicted to be proportional to the number of individuals introduced within natural populations and the life stage of the introduced organisms [21,41]. For instance, in plant populations, it may be more useful to introduce seedlings rather than seeds, as seedlings do not need to meet local germination requirements. Alternatively, introducing seeds may buffer against extreme years and allow germination within more benign conditions [44]. Second, determining the amount and location of introgression of non-local alleles into target population genomes is key for assessing the potential for gene swamping, outbreeding depression, and hybrid vigor. Third, environmental heterogeneity between different target populations or years may or may not create variation in the success of AGF [43]. Both the level of introgression and intensity of selection could fluctuate between years. Finally, evaluating the potential for non-local alleles to move via gene flow between geographically proximate populations is necessary for predicting off target effects (e.g. similar to transgenes into wild populations; [45]). Evaluating the uncertainty and understanding the mechanisms underlying success or failure of AGF provides more confidence for undertaking AGF for threatened or endangered species.

Here, we evaluate short-term fitness consequences after conducting a landscape-scale assisted gene flow experiment in climate-threatened subalpine populations of the common yellow monkeyflower, *Mimulus guttatus* (syn. *Erythranthe guttata*). While *M. guttatus* is widespread across western North America and invasive in the U.K. and New Zealand [46,47], annual populations in the central Oregon Cascades are arguably climate-threatened [44,48,49]. Annual populations of *M. guttatus* are characterized by highly variable growing seasons that depend on ephemeral water availability – growing seasons begin with winter/spring rain or snowmelt and end with terminal droughts [50]. Growing seasons are particular short and variable in annual populations in the central Oregon Cascades, and climate change has both shifted growing seasons earlier and caused more frequent and severe heat waves within these seeps. These populations now exhibit an adaptation lag, where populations from warmer and drier climates with earlier growing seasons (i.e. populations from the foothills of the Sierra Nevada in California) have higher fitness than native Oregon populations [48,51]. Additionally, short heat waves in the last decade have had severe demographic impacts on these populations (50.4% average mortality in one heat wave), with some populations exhibiting near or complete mortality [44,49]. Despite the threatened nature of these particular populations, annual *M. guttatus* continues thrives in other parts of its large range and is not in danger of extinction. These past experiments, along with the substantial genomic resources of this model system, create an ideal scenario to evaluate the impact of assisted gene flow within threatened populations.

We leverage twelve central Oregon Cascade populations with long-term demographic data to conduct our landscape-scale AGF experiment (Fig. 1, Table S1). Populations were clustered geographically into four blocks of three populations with all populations contained within a 25km^2^ region (Blocks A-D below). Baseline data collection began in 2018 with yearly surveys of phenotypes and fecundity in each population. The assisted gene flow experiment began in 2021 with the introduction of individuals from a cross (F_1_) between two maternal lines, one from a low elevation population in the Sierra Nevada (BEL) and one from a high elevation population in the Sierra Nevada (SHL). These populations were chosen because they had high relative fitness in a nearby Oregon common garden over multiple years [44,48,51]. Within each of our experimental blocks, we assigned populations to one of three experimental treatments: control, seeds, or seedlings. In the control populations, we dispersed 200 seeds native to that population. These seeds were pooled from as many different maternal lines as possible from the baseline collections (2018-2020) described above. In the seed treatment populations, we dispersed 200 seeds of the F_1_ source individuals uniformly across the population. In the seedling treatment populations, we planted 19-20 seedlings of the F_1_ source individuals along a transect extending across the center of target populations. We use field surveys and collections before and after assisted gene flow to evaluate the fitness consequences and underlying genetic and phenotypic mechanisms of AGF.

**Fig 1:**
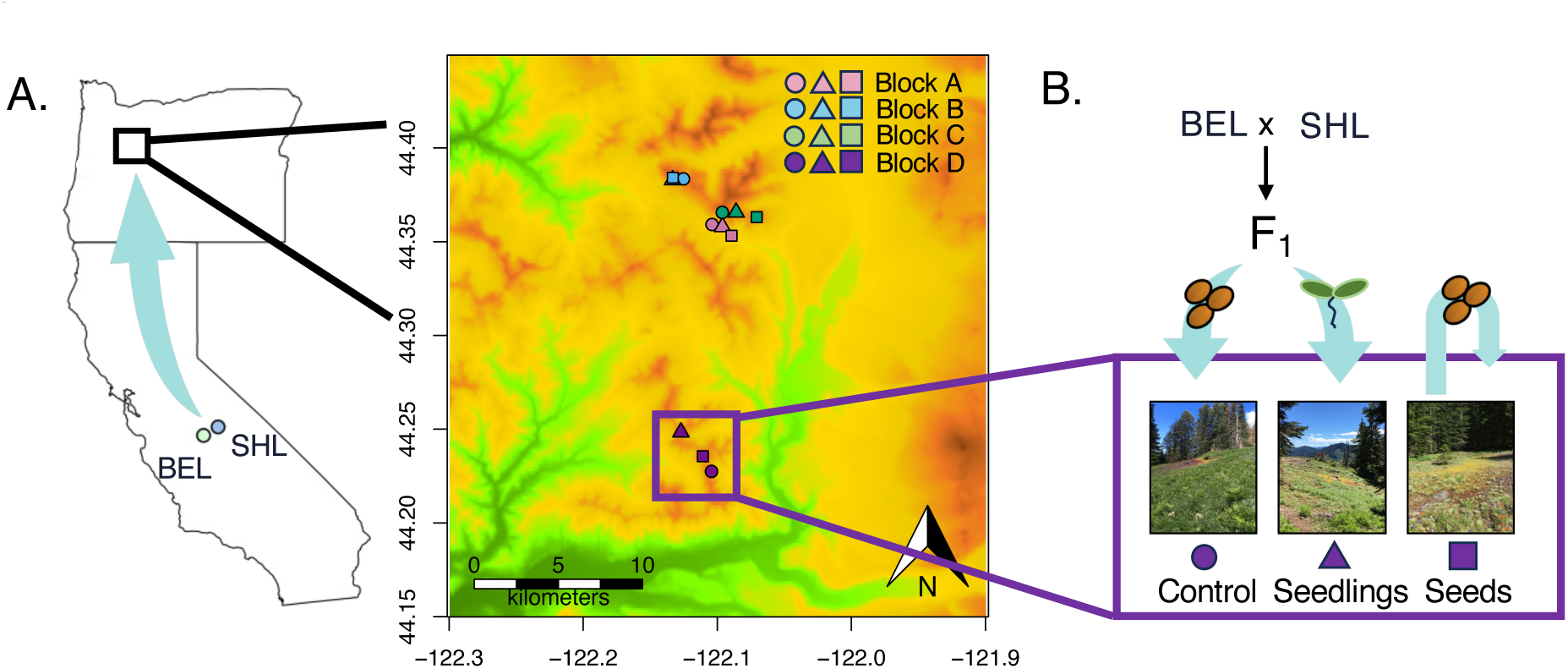
Experimental design and variation in fitness from 2018-2023 across 12 central Oregon Populations. **A.** Map of study area and locations of source populations (BEL and SHL) and target populations for assisted gene flow. Raster depicts elevation with warmer value indicative of higher elevations. Colors represent different experiment blocks and shapes represent treatment (circle = local seeds, triangle = source seedlings, square = source seeds). **B.** Experimental design for assisted gene flow manipulation into each of the four experimental blocks. Each block consists of three populations, one that receives seed from a cross between source populations (F_1_ seed), one that receives seedlings grown from the same cross, and a third that receives seed from the native population. Assisted gene flow occurred in 2021.

## Results

### Fitness consequences of assisted gene flow

Assisted gene flow was associated with modest differences in fitness across the target populations, but the effects differed in direction and magnitude when evaluating different fitness metrics and different life stages of the introduced individuals. Floral abundance increased by 25% post-AGF in populations where California (CA) seeds were introduced and 13.2% post-AGF in populations where CA seedlings were introduced, but only increased by 4.6% in control populations, relative to pre-AGF levels (AGF:treatment, X^2^ = 9.58, p = 0.008; Fig. 2A, Table S2-3). Seed production did not change post-AGF in target populations relative to control populations after conducting assisted gene flow (AGF:treatment, X^2^ = 3.0, p = 0.22). Instead, there were moderate increases in seed production on average across all populations post-AGF, relative to pre-AGF production (28-43%; Fig. 2B). Introduction of novel alleles may also be expected to increase variation in fitness within target populations. Standard deviation in floral abundance increased by 26.4% post-AGF in populations where CA seeds were dispersed and by 3.1% in populations where CA seedlings were planted (Fig. S2A). The standard deviation in seed production decreased by 19.3% in populations where CA seeds were dispersed and increased by 43.0% in populations where CA seedlings were planted (Fig. S2B). These results suggest that assisted gene flow is either weakly effective across all populations or only effective in some of the target populations.

**Fig. 2:**
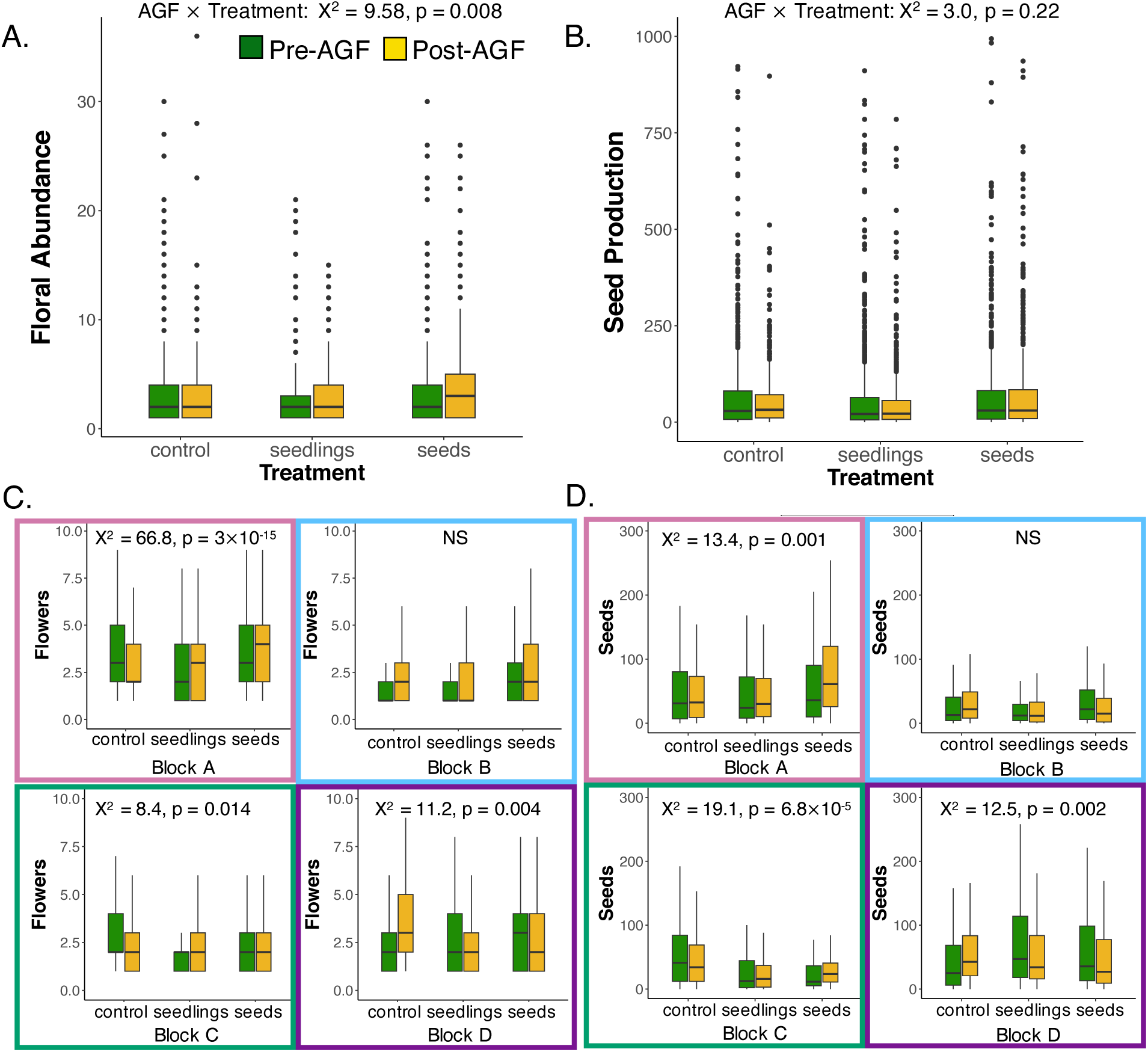
Assessing changes in fitness following assisted gene flow. **A-B**. Boxplots depict changes in floral abundance and seed production following assisted gene flow across treatments: control (native seed introduced), source seeds introduced (CA F_1_), or source seedling introduced (CA F_1_). Each point is an individual. Pre-AGF (green boxes) includes 2018-2021 growing seasons for floral abundance and 2018-2020 growing seasons for seed production. **C-D.** Changes in floral abundance and seed production parsed into the four experimental blocks. Summary statistics and p-values correspond to the AGF:Treatment term within the GLMMs. Border colors on the subpanels correspond to the location of the experimental block on the map in Fig. 1. Edges of boxes in boxplots correspond to the interquartile range, the line inside boxes to medians and the whiskers extend to the most extreme points within 1.5 × IQR from box hinges.

Target populations exhibited substantial heterogeneity in their responses to AGF. Assisted gene flow-related increases in flower abundance were primarily driven by two of the four experimental blocks (Block A and Block C; Fig. 2C, Table S4-5). Conducting AGF via seeds increased floral abundance post-AGF by 62.7% (Block A) and 24.8% (Block C), and conducting AGF via seedlings increased floral abundance by 39.7% (Block A) and 23.7% (Block C) (Block A: AGF:treatment, X^2^ = 66.8, p = 3.1 x 10^-15^; Block C: AGF:treatment, X^2^ = 8.4, p = 0.014). Similar trends were observed in these experimental blocks regarding seed production (Block A: AGF:treatment, X^2^ = 13.4, p = 0.001; Block C: AGF:treatment, X^2^ = 19.1, p = 6.8 x 10^-5^; Fig. 2D). However, AGF resulted in decreases in both floral abundance and seed production post-AGF in a third block relative to control populations (Block D; floral abundance: AGF:treatment, X^2^ = 11.2, p = 0.004, seed production: AGF:treatment, X^2^ = 12.5, p = 0.002). Differences in responses across populations likely depends both on stochastic (e.g., gene flow between plants within populations) and deterministic factors (e.g., seed inviability and the magnitude of early season selection on hybrid individuals).

We observed changes in both the magnitude and direction of fitness responses to AGF in the years following introduction (Fig. S3-S5, Table S6-7). There was a statistically significant increase in floral abundance in AGF treatments in the year following AGF implementation (2022: AGF:treatment, X^2^ = 9.9, p = 0.007) but not in 2023. Seed production exhibited similar heterogeneity between years with greater increases in populations receiving AGF in 2022 and increases in control populations in 2023. There were also differences between populations in the degree of temporal heterogeneity in fitness. For instance, in Block A, introducing CA seeds increased floral abundance most prominently in 2022, but introducing seedlings into the paired population increased floral abundance more prominently in 2023. These results highlight that temporal heterogeneity in selection will have a strong impact on AGF results, but also that some populations exhibit consistent increases in fitness across multiple years.

### Genetic evidence for introgression into AGF-targeted populations

To validate whether the fitness consequences above were consistent with introgression of California alleles into target populations, we grew seeds from each year and population in the UL-Lafayette greenhouse and sequenced whole genomes to an average of 10.9x coverage for 872 individuals. Each population and year combination was represented by 5-27 individuals (Total N = 872; mean = 14.53; SD = 5.2; Table S7). We also sequenced 7 F_1_ lines and 20 plants from the low elevation source population. We examined two lines of evidence with this dataset for evidence of introgression: introgression of private alleles from California and whole genome shifts toward California-like ancestry.

We find evidence of introgression of private alleles into two of the four experimental blocks. In the experimental block with the greatest fitness increases after conducting AGF (Block A), we observed that AGF increased the number of CA private alleles that were successfully incorporated relative to the control population (treatment: X^2^ = 616.7, p = 2.2 x 10^-16^; Fig. 3A, Table S8). Likewise, there is an increase in the number of CA private alleles in the experimental block with the greatest fitness decreases after conducting assisted gene flow (Block D: treatment: X^2^ = 76.151, p = 2.2 x 10^-16^). However, there is no evidence of introgression from private alleles in the other two experiment blocks – including Block C that had increased fitness in AGF treatment relative to control populations. The lack of evidence for genome-wide introgression does not necessarily mean that there were not source alleles introduced, but does indicate that introgression was, at best, rare. In populations with evidence of introgression from source populations, there was no obvious pattern in the distribution of introgression in private alleles across the genome and all private alleles were found at relatively low frequencies in both 2021 and 2022 (Fig. 3B, Fig. S6). Notably, introgression was higher in 2021 than in 2022 and different private alleles introgressed in 2021 and 2022 as only 5.86% of loci have CA alleles present in both years (Table S9). These results confirm that introgression of CA alleles did occur in multiple experimental blocks, and non-local alleles remain at relatively low frequencies in the first years following introduction.

**Fig. 3.**
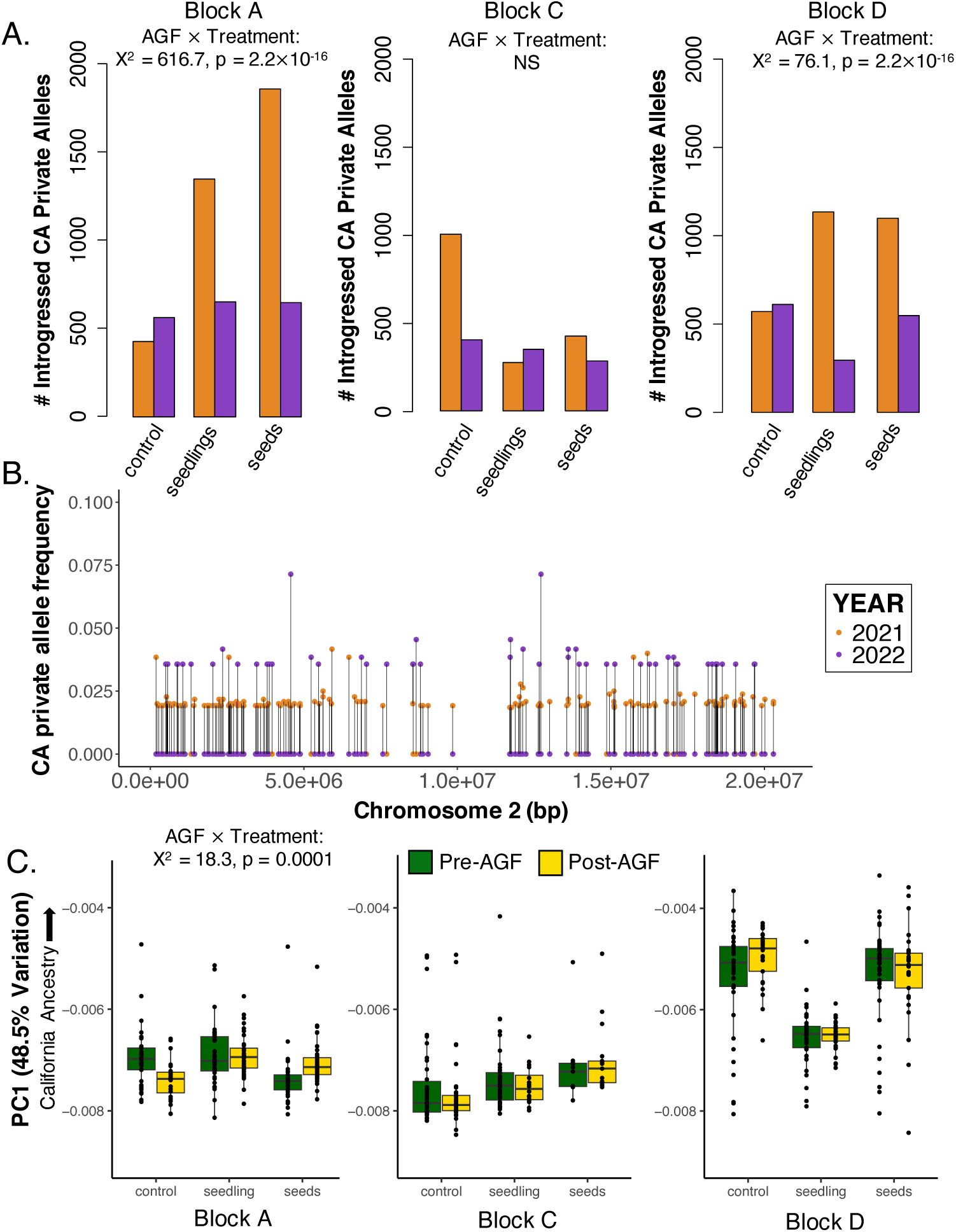
Patterns of introgression into target populations following assisted gene flow. **A**. Number of loci with evidence of introgression of CA private alleles for three experimental blocks. In each subpanel, left, center and right populations correspond to control, seedling and seed population treatments, respectively. Total number of potential loci with California private alleles was 15,691. **B**. Allele frequencies of California private alleles across a representative chromosome (chromosome 2 in the RRM population – a population with source seeds introduced). Allele frequencies may be higher in 2022 because there was greater sample size (27 vs. 14) in 2021. **C.** Variation in genetic principal component 1 before and after assisted gene flow within three experimental blocks. California populations are distinct from Oregon populations with greater values of PC1 (Fig. S7). Edges of boxes in boxplots correspond to the interquartile range, the line inside boxes to medians and the whiskers extend to the most extreme points within 1.5 × IQR from box hinges.

Introgression of CA alleles should also cause AGF target populations to become more genetically similar to CA parents and F_1_ lines (Fig. S7). Genetic principal component analysis conducted with 4-fold degenerate sites suggest that there is substantial genetic divergence between source populations (CA parents/F_1_) and native Oregon populations, represented by PC1 (48.5% of variation). Only one experimental block (Block A) has evidence of a shift toward CA/F_1_ plants along PC1 (AGF:treatment, X^2^ = 18.3, p = 0.0001). Both introducing seeds and introducing seedlings were effective in increasing CA ancestry in this block. However, the magnitude of the shift toward CA ancestry is small and all plants are much closer to Oregon ancestry than California ancestry. This suggests that very low genome-wide levels of introgression can create substantial fitness impacts.

### Phenotype mechanisms underlying AGF fitness responses

Changes in fitness due to the introgression of CA alleles suggests that there should be some underlying change in phenotype that links genetic variation to population performance. We conducted a greenhouse resurrection experiment growing maternal lines collected from all years (2018-2022; Table S10) and populations to determine how phenological, morphological, and physiological phenotypes changed in populations that received AGF relative to control populations. Both flowering time and number of trichomes decreased across populations with introduced CA F_1_ individuals relative to control populations (flowering time: X^2^ = 6.74, p = 0.03; trichomes: X^2^ = 6.8, p = 0.03; Fig. 4AB, Fig. S8-9, Table S11). AGF-driven phenotypic differences were stronger for populations receiving seeds rather than seedlings. Despite this general pattern, there was heterogeneity in phenotype responses across the four experimental blocks. In the population block with evidence of introgression and fitness increases (Block A), flowering time decreased by 6.8 and 3.1 days on average in treatments introducing seeds and seedlings compared to an increase of 0.88 days in the control treatment (Fig. 4C). Trichomes decrease in a similar fashion (Fig. 4D). However, in the experimental block with evidence of introgression and decreased fitness (Block D), flowering time increased by 4.3 and 0.8 days on average in treatments introducing seeds and seedlings (Fig. 4E). Trichomes were not significantly altered in this block (Fig. 4F). These results indicate that phenotypic evolution occurred in the same target populations that had introgression of CA alleles and changes in fitness.

**Fig. 4.**
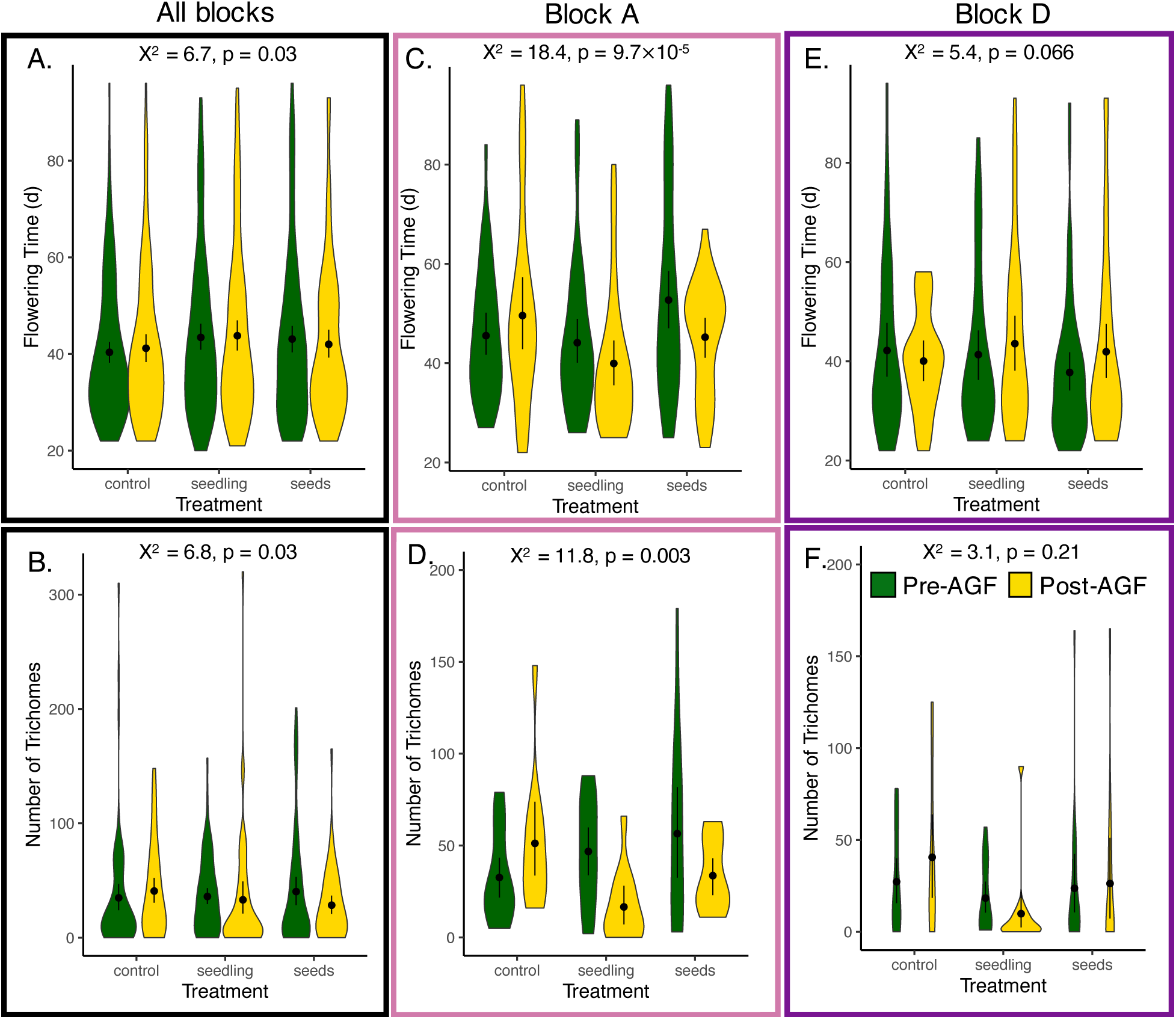
Shifts in flowering time and number of trichomes accompany assisted gene flow. **A-B.** Violin plots depict variation in flowering time and number of trichomes before (green) and after (gold) assisted gene flow. Four populations are pooled within each treatment. **C-F.** Variation in flowering time and number of trichomes from the two experimental blocks with genomic evidence of introgression and fitness impacts of AGF. Fitness was elevated following AFG in block A, but lower in the block D (Fig. 2). Border color of panels corresponds to experiment block color in Fig.1. Points and whiskers inside violin plots correspond to mean and standard error, respectively. Summary statistics and p-values correspond to the AGF:Treatment term within the GLMMs.

We hypothesize that phenotypic evolution in AGF-target population populations should be in the direction of source California populations. Previous greenhouse common garden studies suggest both California parent populations flower more quickly on average than any of our 12 central Oregon focal populations (30.7-35 mean days following germination; Kooyers et al. 2015) and have high levels of variation in trichome production (range: 6-250 trichomes; Kooyers et al. 2015). There are individuals in every focal Oregon population that flower as quickly as California populations. However, populations in Block A were 3 of the 4 slowest flowering central Oregon populations prior to AGF and evolve earlier flowering in the direction of the California parent lines. Populations in block D, particularly BMP, had relatively faster times to flowering prior to AGF that were more closely resembled California source populations. Thus, evolution of later flowering within this block would not have been predicted, but could be generated if introgression broke up co-adapted combinations of alleles. We cannot make the same predictions for the evolution in trichome number as the source populations are too variable. Taken together, these results suggest that the introgression of California alleles altered flowering time and created variation in fitness following AGF.

## Discussion

We find that assisted gene flow has the potential to increase fitness of target populations within only a few generations. Theoretical models simulating genetic or evolutionary rescue indicate that introduction of foreign alleles often lead to initial maladaptation due to outbreeding depression followed by increases in fitness as adaptive alleles are selected and maladaptive alleles purged from the population [36,37]. We find two blocks that had evidence of introgression and that also exhibited changes in fitness post AGF. Both target populations in one block with introgression exhibited slight evidence of maladaptation or outbreeding depression relative to control populations in the first three generations, consistent with theoretical expectations (Fig. 2). Surveys over the next decade will determine whether introgression eventually increases population fitness in this initially maladapted block. However, target populations in the other experimental block with introgression had substantial fitness gains within these first generations (Fig. 2). There are multiple non-mutually exclusive hypotheses explaining this rapid fitness increase. One explanation for fitness gains is hybrid vigor or heterosis, where outbreeding leads to increased heterozygosity and an increase in fitness [52]. However, since allele frequencies associated with the introgression of private alleles are low (Fig. 3), this is unlikely to be the driving mechanism. Also unlikely is a scenario where the influx of genetic diversity rescues the population from inbreeding or high genetic load as target populations have high levels of genetic diversity [53]. The most likely possibility is that target populations were poorly adapted to their local conditions and AGF introduced novel genetic variants that shifted phenotype means and allowed transgressive phenotypes or novel phenotypic combinations to emerge. While our greenhouse work establishes post-AGF shifts toward earlier flowering and lower trichomes relative to control populations, field common gardens with manipulations to key climate variables (e.g. growing season length) are necessary to dissect whether the observed phenotypic evolution could underlie our observed fitness gains.

One potential caveat to our results is that fitness differences pre- and post-AGF could be associated with intrinsic differences between the individual populations within a block, rather the AGF treatments. Indeed, if a control population were to have declining fitness following AGF due to any random reason, both treatment populations would appear to have enhanced fitness. We believe our results are robust, particularly for increase in fitness associated with AGF in Block A as well as the decrease in fitness in Block D, for three reasons. First, we control for intrinsic differences among population (and years) within our models using random effects. Second, populations *within* blocks are geographically paired and nearly indistinguishable in population structure analysis (with the exception of low levels of differentiation in Block D). Finally, the same target populations that exhibit fitness differences in certain years also exhibit phenotypic evolution and show the strongest signals of introgression (Block A &D). Thus, although our results are necessarily correlative because of our landscape-scale experimental design, these results strongly suggest that AGF caused introgression, produced phenotypic evolution, and altered fitness of focal populations.

### Climatic variation impacts introgression and fitness gains associated with AGF

Our results provide key insights into the expectations for assisted gene flow within complex environments. We find variation in the success of assisted gene flow across both space and time (Fig. 2, Fig S4). Our finding that two experimental blocks had evidence of introgression from assisted gene flow while two other blocks did not could be explained by both deterministic and stochastic factors. Lack of introgression could stem from stochastic factors. For instance, relatively few seeds or seedlings were introduced and introgression is dependent on pollination between native and introduced individuals.

However, success of introgression and associated fitness differences were likely impacted by the climatic differences between populations. While populations within each experimental block were located at the same elevation, experimental blocks were located over an elevation gradient of ∼500m – creating substantial differences in growing season timing and both abiotic and biotic conditions during the growing season. The lowest elevation experimental block was the block with evidence of introgression, phenotypic evolution and fitness gains. These populations have an earlier growing season start date and a longer growth season than higher elevation populations – conditions that are most similar to the historical conditions with the source California populations [50]. Notably, low elevation populations also had more delayed times to flowering than high elevation populations at the beginning of the experiment. An infusion of novel alleles promoting earlier flowering, such as those from the early flowering California source plants, would be most likely to be effective in these populations and could be detrimental in target populations that already flower earlier than the source populations, i.e., are already better adapted to changing conditions. Conditions for germination of seeds may also have differed between populations and this is likely a key barrier for the success of assisted gene flow [54]. Our results suggest, broadly, that the success of AGF is dependent on whether local populations are already well adapted to the environmental conditions during the translocation years.

The difference in introgression and fitness in different years following AGF also suggests that climate abnormalities in the initial years of AGF act as an important filter for AGF success. Patterns of introgression differ substantially between years. Most introgression took place during the year that seeds and seedling were introduced (2021). This should be expected as seeds germinate readily in *M. guttatus* and we planted a number of seedlings that were ready to flower (Fig. S1). However, there was also a substantial heat dome event during the 2021 growing season [55] that had devastating consequences on many target populations [49] and prevented many of the source seedlings from producing seed (Table S12). We also expected evidence of introgression in subsequent years from seeds that did not germinate the first year and from seeds which were generated as hybrids between source and target populations. There was far more limited evidence of introgression of private alleles relative to control populations in 2022 (Fig. 3). It is notable that we judge levels of introgression relative to control populations as the likely pollinators (bees) are capable of flying between control and treatment populations within an experimental block (1-2km; 55). Indeed, alleles that we inferred as private to source populations appear in every control population (Fig. 3). This is primarily due to our inference of private alleles as we likely did not detect some rare alleles in Oregon in the sequencing of 2018-20 Oregon populations. However, tracking source population alleles that escape treatment populations is important for determining off-target impacts of AGF [57].

Limited introgression of private alleles genome-wide does not exclude the possibility that there may be introgression in key genomic regions that could have impacted phenotypes. On the contrary, some populations with relatively limited introgression of private alleles in 2022 had strongly increased floral abundance relative to control populations in the same year (e.g. Block A seeds treatment). Future studies scanning the genome for allele frequency change will detect the key regions underlying shifts in fitness. However, temporal variance in relative fitness in AGF populations relative to control populations likely indicates that temporal variation in climate plays a key role in the outcome of AGF. High elevation *M. guttatus* populations in the central Oregon Cascades are known for strong fluctuating selection maintaining high genetic and phenotypic diversity within populations [53,58–60]. Notably, one of the key phenotypes undergoing fluctuating selection is flowering time with year-to-year variation driven by differences in growing season start date and duration [59]. An important consideration for AGF is that introduced alleles may be adaptive in some years but not in others. For instance, we worry that introduced alleles that increase in frequency during the relatively benign years (i.e. 2022 growing season) may be maladaptive and cause populations to be less resilient during future heatwaves. Moreover, strong fluctuations during the first years after AGF could either enhance or prevent source population alleles from incorporating into target populations depending on whether the fluctuation makes the environment more similar to the historic source environment. The relative impacts of fluctuating selection during AGF require additional theoretical work.

### Assessing key choices for implementing AGF

Our results suggest that the logistics of assisted gene flow are key to the end result. Current recommendations suggest introducing 5-20% of the target population census size from source populations [21,41]. Our experiment introduced ∼ 0.5 - 2 % of the census size depending on the population and whether we introduced seeds or seedlings. Population census sizes can differ dramatically between years, with limited individuals flowering during years with early growing season start dates and abnormally hot conditions [44,49]. This decision likely limited the initial amount of introgression in target populations. Notably, even with a limited number of individuals introduced, we observed both introgression and fitness impacts within half of our target populations. Despite this success, we suggest introducing higher numbers of individuals to ensure more rapid incorporation into the genome (i.e., 10-20% of census size).

Another key choice is deciding how to introduce source populations. Our experiment examines a single introduction of seeds vs. seedlings. In experimental blocks with clear evidence of introgression, both seed and seedling treatments had effects, but the introduction of seeds increased introgression and led to larger magnitude fitness effects. The introduction of seeds was likely more effective for three reasons: first, we introduced more seeds than planted seedlings; second, 77% of seedlings died following introduction due to a substantial heatwave and the rest produced limited fruits (1.17 ± 0.99 SD fruits); third, seeds may not have germinated within the first year and could have provided a reservoir capable of introducing alleles in multiple growing seasons. Thus, we do not observe the potential downside of introducing seeds – that they must overcome seed dormancy requirements and novel establishment conditions associated with non-native sites [54]. Our results suggest that single introductions are less likely to be effective than multiple introductions and that introducing seeds may have longer lasting effects in unpredictable climates.

Our experiment also included a non-standard choice for applied assisted gene flow efforts. We introduced germplasm resulting from a cross of two maternal lines sourced from two populations. Our choice was justified by the success of these two lines and populations in a nearby common garden experiment [44,48], evidence of inbreeding depression in nearby natural populations [61], and the need for a genetically-tractable system. Yet, most species with the need for AGF will not have the large history of evolutionary ecology and genetics studies that our study system possesses. Source populations for AGF are typically selected via climate transfer zones or seedlot selection tools based on historical climates and contemporary conditions [62]. Experimental and simulation studies are needed to better assess how to choose source populations – phenotypic and genomic data could provide rapid assessment of potential source populations that could complement or even replace climate data. In the absence of data, including both native and climate-adapted germplasm in introductions either via seed mixes or through hybridization of populations can allow natural selection to incorporate ecologically-important variants.

### Conclusions

Our study provides cautious support for utilizing assisted gene flow to preserve natural populations experiencing the detrimental effects of climate change, consistent with the growing recommendations for conducting AGF in a variety of maladapted populations [16,17,63,64]. We find evidence of the beginning stages of evolutionary rescue despite relatively limited genome-wide patterns of introgression, limiting potential concerns regarding outbreeding depression or gene swamping. However, attention to organismal biology of the threatened species and logistical details is clearly key to the success or failure of AGF and we encourage responsible evidence-based decision making.

## Materials and Methods

We conducted a landscape-scale manipulative experiment involving 12 target populations within a 25km^2^ region of the Central Oregon Cascade Mountains. Populations were geographically clustered into four groups of three populations (‘Blocks’) with 0.5 - 2km between populations within blocks (Fig. 1). Elevations were similar within blocks but varied between blocks (1087-1567 m; Table S1). We selected two source populations from the foothills of the Sierra Nevada in central California (Fig. 1; Table S1). These populations were chosen because they had higher fitness than the native populations from the Oregon Cascades in nearby common garden experiments [48,65]. A single maternal line from the two source populations (BEL, SHL) were crossed together in bulk to produce the F_1_ seeds used for AGF. In each block, populations were designated as control, seed or seedling treatment. In the control treatment populations, 200 seeds previously collected from the native population were uniformly sown throughout the population (20+ maternal lines). In the seed treatment populations, we dispersed 200 F_1_ source seeds uniformly across the population. In the seedling treatment populations, we planted 19-20 14-day old seedlings grown from the F_1_ seeds along a transect extending across the center of target populations. The difference in sample size reflects germination and establishment likely provide a substantial demographic filter within these populations. Treatments occurred as soon as possible after snowmelt to correspond to population phenology. For seedling growth conditions prior to planting, see extended methods. We recorded survival of translocated seedlings weekly, and counted number of fruits produced after senescence in order to assess the potential for introgression of new alleles in these populations; only 23% of seedlings survived to flower, and those individuals produced an average of 1.17 ± 0.99 SD fruits (Table S11). However, we do note that a single fruit can contain hundreds of seeds, and natural populations also had very few individuals surviving to reproduction and producing seeds during these years [49].

### Evaluating AGF impacts on fitness within natural populations

We assessed fitness at the end of the growing season in each population in each year from 2018-2023 (previously reported in [44,49]. Plants were collected along two 7.5 m transects spanning the population. From 2018-2020, one individual was collected every 15 cm along each transect, totaling 100 individuals/population. From 2021-2023, one individual was collected every 30 cm along each transect, totaling 50 individuals/population. We counted all flowers and seeds for each individual as proxies of fitness. We note that assessing population demography is methodically challenging within this species. Central Oregon populations exhibit sweepstakes reproduction where an individual seed has a low chance of producing seeds in the next generation. Specifically, seeds must land in ephemerally wet patches within an extremely heterogeneous seepy meadow. Thus, fecundity (i.e., increasing the number of chances at reproductive success) is likely strongly associated with population growth rate, but our work cannot assess whether AGF increases population growth rates above replacement level.

We evaluated whether AGF increased fecundity of the target populations within a general linear mixed model framework (LMM). We conducted univariate LMM to assess the effects of AGF (pre/post), Treatment (control/seeds/seedlings), and the interaction between AGF:Treatment on both metrics of fitness (flower abundance and seed production). Year and population were random effects in each model. Floral abundance and seed production were log+1-transformed to better fit model assumptions. To determine whether assisted gene flow increases the variance in fitness, we conduced similar univariate LMMs (without population as a random factor) using the standard deviation of floral abundance and seed production between individuals within a population as response variables. Standard deviation of flower abundance was log+1-transformed. Analyses were performed using the *lme4* v 1.1-35.5 package in R v 4.4.1 [66]. Statistical significance of LMM was assessed via ANOVA with type III sum of squares using the Anova() function in the *car* v 3.1-2 package [67]. We note that interpretation of models differed for each fitness variable as flower abundances could logistically not be affected by AGF in 2021 as plants sampled along transects were almost certainly native Oregon plants (i.e., not from seed germinated from source populations); thus, floral abundance in 2021 is considered pre-AGF in models. However, seed production could have been impacted in 2021 through pollen transfer from introduced CA seeds/seedlings and can be considered post-AGF in models.

To better dissect the impacts of AGF in the experimental manipulation, we evaluated variation among blocks and between post-AGF years. We examined differences among different experimental blocks by subsetting the data into the four blocks and repeating the above univariate LMMs for each block. Because each treatment was only represented by one population, population was removed as a random factor for these models. We assessed heterogeneity in AGF-related fitness variation across post-AGF years by subsetting data into each post-AGF year and retaining all baseline years. In these models, year was removed and population was included as a random variable.

### Validating AGF introgression via genomic analysis

We germinated seeds from 24-30 maternal lines/population ∼2 months after collection each year in the UL-Lafayette Greenhouse (*Refresher Generation Growth Conditions* in extended methods). We also included 7 individuals from one parental population (BEL) and 20 F1 individuals grown in the 2021 common garden. Seeds from the other parent line (SHL) did not germinate. One individual from each maternal line had buds or leaves collected for genetics and this individual was selfed for use in the resurrection experiment described below. DNA was extracted using a modified CTAB extraction [68] and diluted to 3 ng/μL for library construction. Libraries for whole genome sequencing were constructed using a customized dual-index Nextera library construction protocol [69]. Each population and year combination was represented by 5-27 individuals (Total N = 872; mean = 14.53; SD = 5.2; Table S7). Poor fitness in the field that limited collections as well as reduced germination led to the reduced sample size of some populations in some years. Post-barcoding, we pooled either 48 or 96 individual equimolar libraries and sequenced each pool on a single lane on either a HiSeqX or NovoseqX Plus platforms (150bp paired end reads; Novogene, Davis, CA). Batch effects due to either sequencing platform or lane were minimal.

We constructed a custom bioinformatics pipeline for our population genomics study (https://github.com/donnasaurrus/AGF). Initial read quality was evaluated using fastQC [70]. High-quality reads were aligned a *Mimulus guttatus* reference genome from a nearby population (IM62 v3: https://phytozome-next.jgi.doe.gov/info/Mguttatusvar_IM62_v3_1) utilizing bwa-mem with default parameters [71] and converted to bam files using SAMtools [72]. Read groups and duplicates were marked using PicardTools, and SNPs were called using bcftools [73]. We filtered for variant quality, strand bias, and mapping quality using GATK [QD < 2.0 || FS > 60.0 || MQ < 40.0 || MQRankSum < -12.5] [74]. We used VCFtools [75] to filter retaining biallelic SNPs with minimum quality score of 500, and removing any SNPs with >25% missing data. This SNP dataset included 18,118,076 SNPs. Mean coverage and error rates of our sequence data (.bam files) were assessed using Qualimap [76,77]. Mean coverage depth was 10.9x (SD = 7.7), and mean error rate was 2.7% (SD = 0.39%) (Table S13).

We first evaluated introgression via AGF into Oregon populations using private alleles. We filtered the SNP dataset to only include sites where both reference and alternative alleles were at a frequency > 0.3 in BEL and F_1_ individuals *and* were monomorphic in all populations in Oregon prior to conducting AGF (2018-2020). We note that these alleles may not necessarily be truly private as some rare alleles in Oregon populations may escape detection in the three baseline years, either due to the presence of a seed bank or a finite sample size. This dataset consisted of 15,691 SNPs distributed genome-wide. We evaluated whether there was introgression in target populations that received AGF from source populations compared to control populations. We transformed allele frequencies into a binomial variable, where loci with a CA private allele frequency of 0 were coded as 0, and any non-zero allele frequencies were coded as 1. We conducted a generalized linear mixed model with a binomial distribution and a logit link function using the *glmmTMB* v 1.1.9 package in R v 4.4.1 [78], with treatment (levels: Control, Seed, Seedling) included as a fixed factor, and with year and population included as random variables. To assess differential responses between blocks, we repeated this model, subsetting data into individual experimental blocks and removing population as a random variable. Statistical significance was assessed as in above GLMs for fitness.

We evaluated the extent that introgression shifted genome-wide genetic diversity of Oregon populations toward California source populations using a principal component analysis that included Oregon populations from all years as well as individuals from California parental and F_1_ lines. We filtered SNPs to include only 4-fold degenerate sites and only used individuals with < 20% missing data (N = 803). This dataset included 1,056,971 SNPs. A PCA was conducted using --pca in *plink* v1.9 [79]. PC1 separated Oregon from California populations (Fig. S7). We examined shifts in PC1 toward CA populations following AGF using univariate GLMs identical those used above for fitness except with PC1 and PC2 as response variables. In these models, AGF (pre/post), treatment (control/seeds/seedlings), and the interaction between AGF:Tntroduction were fixed effects while population and year were included as random effects.

### Identifying phenotypic shifts following AGF

We conducted a greenhouse resurrection experiment to examine how phenotypes shift following AGF. We germinated seed from 988 maternal lines stemming from every population and year from 2018 - 2022 (mean ± SD: 17.0 ± 6.1 lines per population per year, Table S9). These seeds were selfed lines from the refresher generation(s) described above. Germination and growth conditions are described in extended methods (*Resurrection Experiment Growth Conditions*). We surveyed for germination and flowering daily and calculated flowering time as the number of days between germination and opening of the first flower. At flowering, we identified the node of the first flower, counted the number of leaves and branches, and measured plant height, corolla width, corolla length, corolla height, and stem diameter. We also collected the larger single 2^nd^ true leaf to measure leaf area, specific leaf area, number of trichomes, and relative water content. Further details in phenotypes can be found in extended methods. There were strong correlations among several of the phenotypes (Table S14) so we choose to only include a subset of phenotypes in analyses including flowering time, plant height, leaf number, corolla width and number of trichomes.

We used univariate GLMMs to evaluate potential shifts in phenotypes following AGF treatment. As in above GLMMs, AGF (pre/post), Treatment (control/seeds/seedlings), and the interaction between AGF:Treatment were fixed effects while population and year were included as random effects. Distributions, links, and transformations were chosen to maximize model fit. The GLMM for flowering time had a Poisson distribution with a log link. All other variables were log-transformed and had Gaussian distributions with identity links. Statistical significance of fixed factors was assessed via ANOVA with type III sum of squares using the Anova() function in the *car* v3.1-2 R package [67]. Changes in phenotype due to AGF would create significant AGF:Treatment interactions with shifts likely toward CA phenotypic means. To better examine how phenotypic change was occurring within experimental blocks, we subset the dataset by block and ran identical models for each block without population as a random factor.

## Acknowledgements

The authors thank Josh FitzPatrick, Braden Doucet, Luke Rabalais, Elissa Harb, Jace Segura, Gabbie Dietel, Lana Gaspard, Noah Richard, Emily Bollich, Haley St. Martin, Laura McDonald, Anna Scharnagl, and Benjamin LeBlanc who helped with field, greenhouse and lab work associated with this project. This work was logistically supported by H.J. Andrews Experimental Forest and permitted through the Willamette Forest District of the United States Forest Service. Funding for this research came from University of Louisiana, Lafayette and NSF grants DEB-2045643 and IOS-2222466 to N.J.K.

## Author Contributions

N.J.K. conceived and designed the experiments. DMH, CMP, AT, SDH, SGI, and NJK collected data and performed the experiments. DMH and NJK analyzed the data. DMH and NJK wrote the original draft of the manuscript, and all authors reviewed and edited the manuscript. All authors approved of the final submission.

## Data Availability Statement

Datasets containing field fitness data and phenotypic data from the resurrection experiment will be deposited on Dryad. Sequence data will be uploaded to NCBI SRA as .fastq files.

## Extended Data

### Extended Methods

**Fig. S1:** Seedling transplant into a central Oregon Population (FIR) at different developmental stages.

**Fig. S2:** Changes in variation in fitness following assisted gene flow.

**Fig. S3:** Variation in floral abundance and seed production across six growing seasons and twelve populations

**Fig. S4:** Relative changes in floral abundance and seed production across years following assisted gene flow.

**Fig. S5:** Temporal variation in fitness during the assisted gene flow experiment.

**Fig. S6:** Limited introgression of California private alleles within target populations.

**Fig. S7:** Populations structure across source and target monkeyflower populations.

**Fig. S8:** Variation between pre- and post-assisted gene flow in flowering time within a common garden by experimental block

**Fig. S9:** Variation between pre- and post-assisted gene flow in number of trichomes within a common garden by experimental block.

### Supplemental Tables

**Table S1**: Population geographic locations and treatments

**Table S2:** Model summaries of fitness effects of assisted gene flow within Oregon populations

**Table S3:** Estimated marginal means and raw means for of fitness effects of assisted gene flow across Oregon populations

**Table S4:** ANOVA summary of fitness effects of assisted gene flow within each block

**Table S5:** Estimated marginal means and raw means for of fitness effects of assisted gene flow within individual blocks

**Table S6:** ANOVA summary of fitness effects in individual years following assisted gene flow

**Table S7:** Sampling summary for genomic analysis

**Table S8:** Generalized linear model summaries for private allele analysis

**Table S9:** Counts of private alleles across target Oregon populations and years

**Table S10:** Sampling summary from the resurrection experiment

**Table S11:** Generalized linear modeling results from the resurrection experiment

**Table S12:** Source seedling performance in target Oregon populations

**Table S13**: Population summary of coverage and mean depth for genomic analysis

**Table S14:** Phenotypic correlations in the resurrection experiment

