## Extended Methods for "Rapid adaptation follows experimental assisted gene flow in subset of annual monkeyflower populations"

*Seedling Growth Conditions Prior to Transplant*

$F_1$ seedlings from California source populations were grown in the University of Oregon greenhouse concurrent with the timing of germination in natural populations. We planted $F_1$ seeds into 2.25" square pots filled with Fafard 3B potting soil. Pots were randomized into 1020 flats that were bottom-watered and covered with humidity domes for a 7 day cold-stratification treatment at 4°C in a dark cold room at University of Oregon. Flats were moved to the University of Oregon Greenhouse under ambient light and temperature conditions. We removed humidity domes after 7 days in the greenhouse. We planted seedlings in the field after 14 d in the greenhouse. Seedlings had 1-2 true leaf pairs at planting, and effort was made to plant seedlings as plugs to avoid catastrophic root damage and improve establishment.

*Refresher Generation Growth Conditions*

Each year, following collection of seeds and fitness data from maternal lines in the field, we conducted a common garden 'refresher' generation in UL-Lafayette Greenhouse to remove maternal effects. These growth conditions and experiments were already described in Kooyers et al. 2025. We planted seeds from 24-30 maternal lines per population per year in 2.25" square pots with Sunshine Metro Mix 830 Soil. Less lines or seeds per pot were planted from populations or years that had limited seed production in the field (i.e., HDM was extirpated from 2019-2021 and had limited number of individuals following extirpation). Pots were organized into 1020 flats, covered with humidity domes and stratified for 7 days at 4°C in a dark cold room at University of Louisiana. Flats were then moved to growth shelving with four growth lights (1.22-m 8-bulb T5 fluorescent fixture) with timers set for 16 h day/8 h night cycles at 23°C. We surveyed germination daily and thinned pots to a single germinant closest to the center of each pot. After 7 days, flats were moved to greenhouse and humidity domes were removed. Greenhouses were set at 22°C with supplemental lighting (16 h day/8 h night). At maturity, each plant was self-pollinated and seeds were collected for the resurrection experiment described below.

*Resurrection Experiment Growth Conditions*

We conducted the resurrection experiment in the fall of 2023 in the UL-Lafayette greenhouse under the same growth conditions as in the refresher generation described above. Seeds from selfed lines from the refresher generations (2018-2022) were planted in 2.25" square pots in Sunshine Metro Mix 830 media and moved into 1020 flats with humidity domes. Pots were cold-stratified at 4°C in the dark for seven days. Germination occurred in the greenhouse where natural sunlight was supplemented with grow lights (1.22-m 8-bulb T5 fluorescent fixture) set to a 16h/8h day/night cycle at 22°C. Flats were misted daily for the first seven days, bottom-watered as needed, and rotated daily. Humidity domes were removed after seven days, pots were thinned to a single plant, and randomized within flats (N = 578 total plants). Germination was recorded daily for the first 10 days. Greenhouse conditions included 16h/8h day/night light cycles with a temperature set to 22°C.

*Resurrection Experiment Phenotypic Details.*

We surveyed morphological, phenological, and physiological phenotypes in the resurrection experiment. We checked plants daily for flowering and calculated flowering time as the number of days between germination and the opening of the first flower. Several phenotypes were recorded at the date of first flowering including node of the first flower, number of leaves, number of branches, plant height, corolla width, corolla length, corolla height and stem diameter. Number of leaves excluded the cotyledons. Plant height was measured as the distance from the top of the apical meristem to the stem:soil contact point. Stem diameter was measured directly beneath the 2$^{nd}$ leaf node. We also collected the larger 2$^{nd}$ true leaf to measure number of trichomes, leaf area, specific leaf area, and relative water content. Following excision, leaves were floated, peduncle down, in deionized water for 12-16hrs. Saturated leaves were weighed ('wet leaf mass') and photographs were taken to calculate leaf area. Leaf area was assessed in imageJ relative to a 1cm$^2$ square. We counted number of trichomes under a dissecting microscope; we counted all trichomes along a lateral transect of the widest section of the leaf. Leaves were dried at 65°C for at least 4 days – a time period sufficient to remove the maximum amount of water from the leaf – before weighing each leaf ('dry leaf mass'). We calculated specific leaf area as leaf area / dry leaf mass and calculated relative water content as (wet leaf mass – dry leaf mass) / wet leaf mass.
