## Supplemental Figures for "Rapid adaptation follows experimental assisted gene flow in subset of annual monkeyflower populations"

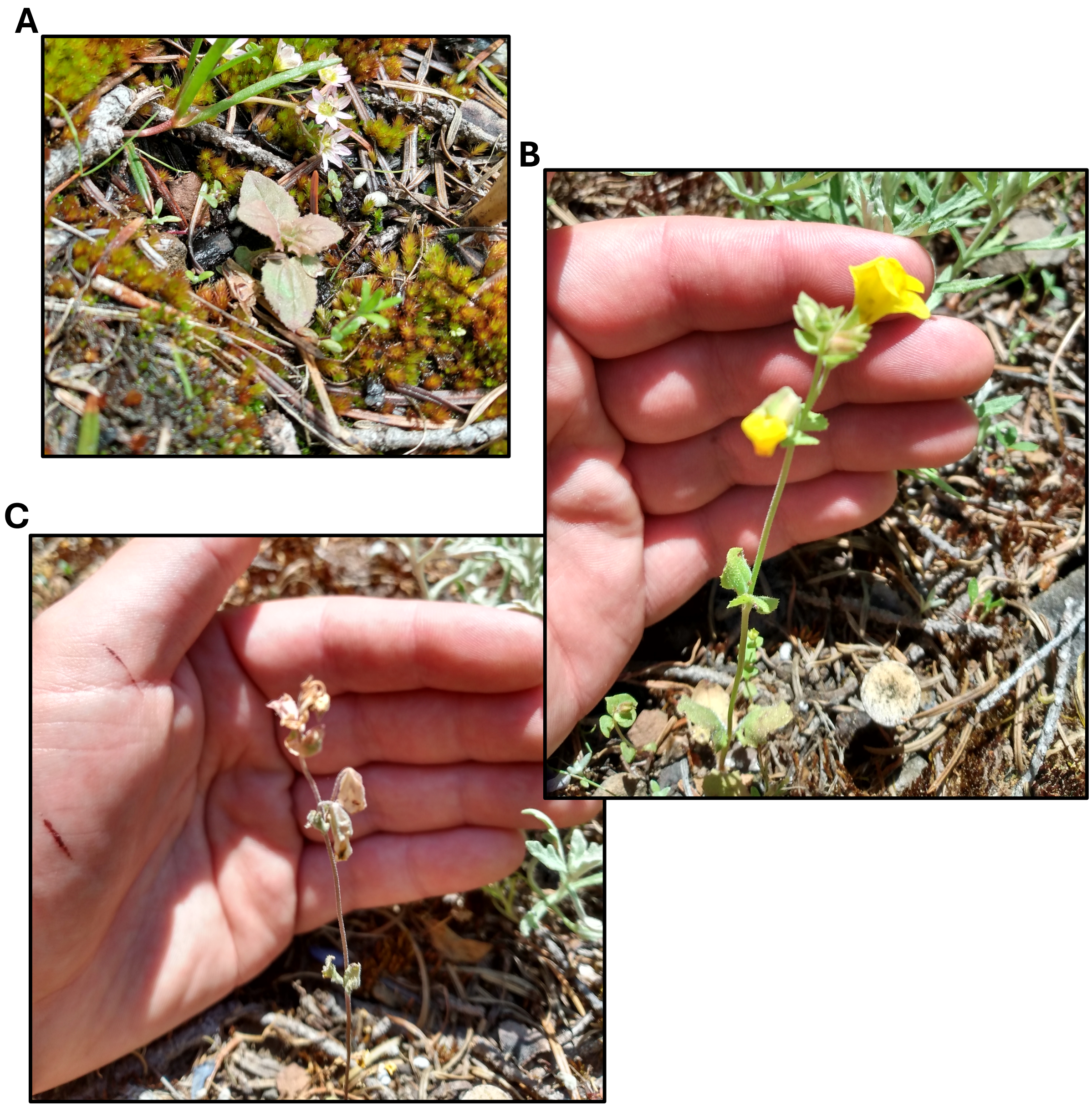


**Fig. S1: Seedling transplant into a central Oregon Population (FIR) at different developmental stages. A.** One week after transplanting. **B.** At time of first flower. **C.** After senescence.

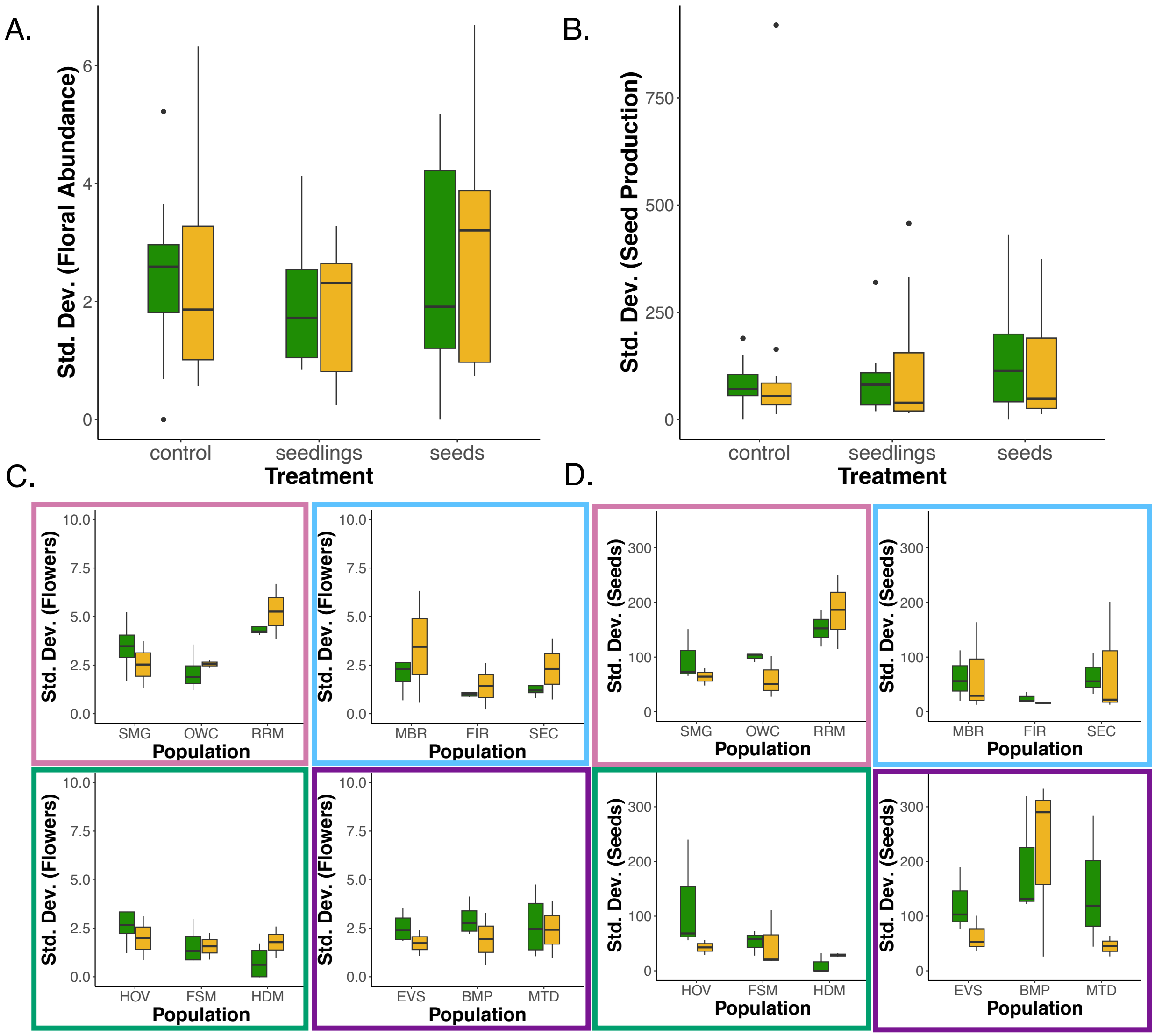


**Fig. S2: Changes in variation in fitness following assisted gene flow**. **A-B**. Boxplots depict changes in the standard deviation in floral abundance and seed production following assisted gene flow across treatments: control (native seed introduced), source seeds introduced (CA $F_1$), or source seedlings introduced (CA $F_1$). Each point is an individual. Pre-AGF (green boxes) includes 2018-2021 growing seasons for floral abundance and 2018-2020 growing seasons for seed production. **C-D.** Shifts in the standard deviation of floral abundance and seed production parsed into the four experimental blocks. In each subpanel, left, center and right populations correspond to control, seedlings and seed population treatments, respectively. Border colors on the subpanels correspond to the location of the experimental block on the map in Fig. 1. Edges of boxes in boxplots correspond to the interquartile range, the line inside boxes to medians and the whiskers extend to the most extreme points within 1.5 × IQR from box hinges.

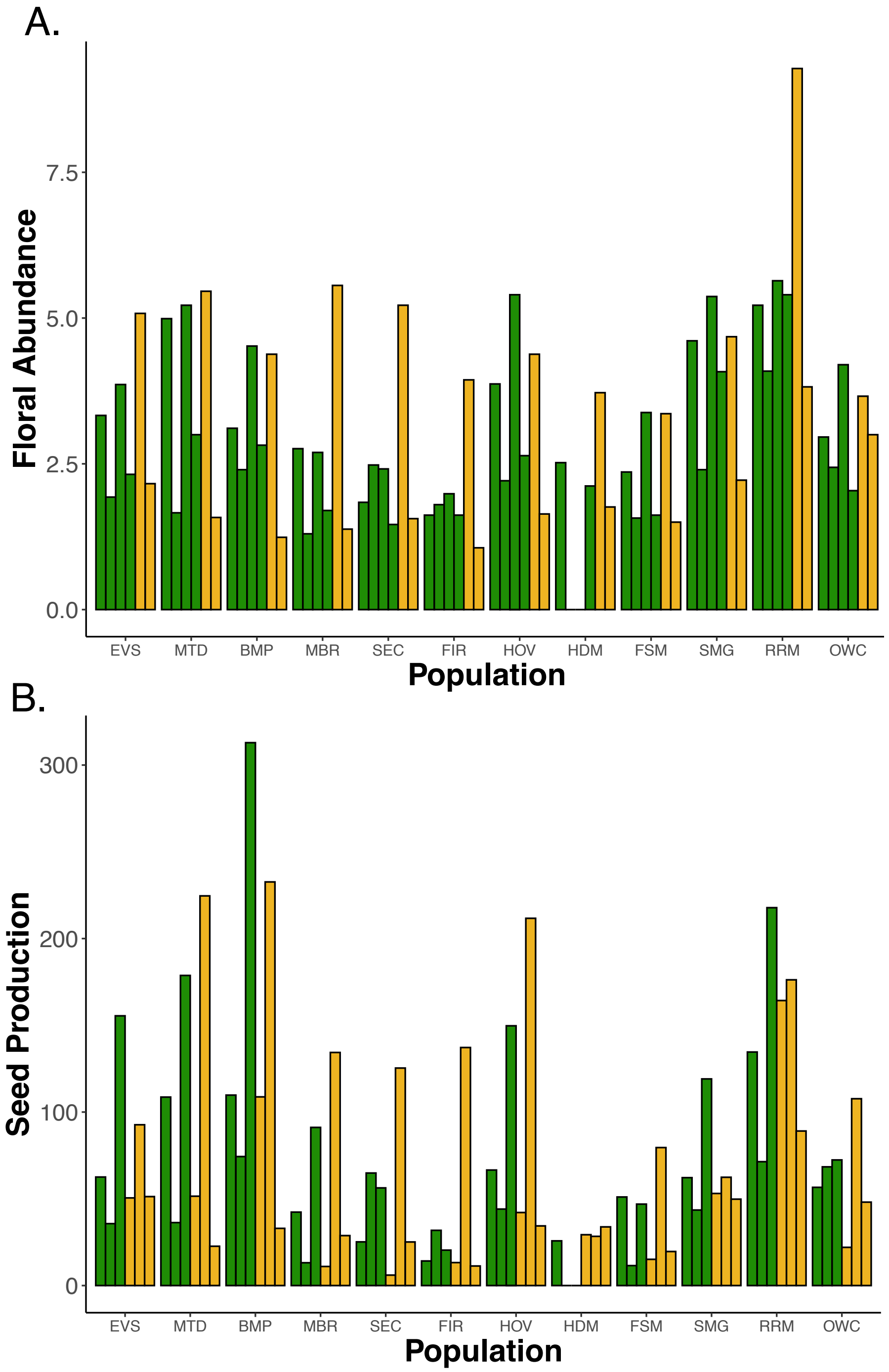


**Fig. S3: Variation in floral abundance and seed production across six growing seasons and twelve populations**. Green bars correspond to years before assisted gene flow and yellow bars correspond to years after assisted gene flow. Years are in order (2018-2023). EVS, MBR, HOV and SMG are control populations. Source seeds were introduced to MTD, SEC, HDM and RRM. Source seedlings were introduced to BMP, FIR, FSM, and OWC.

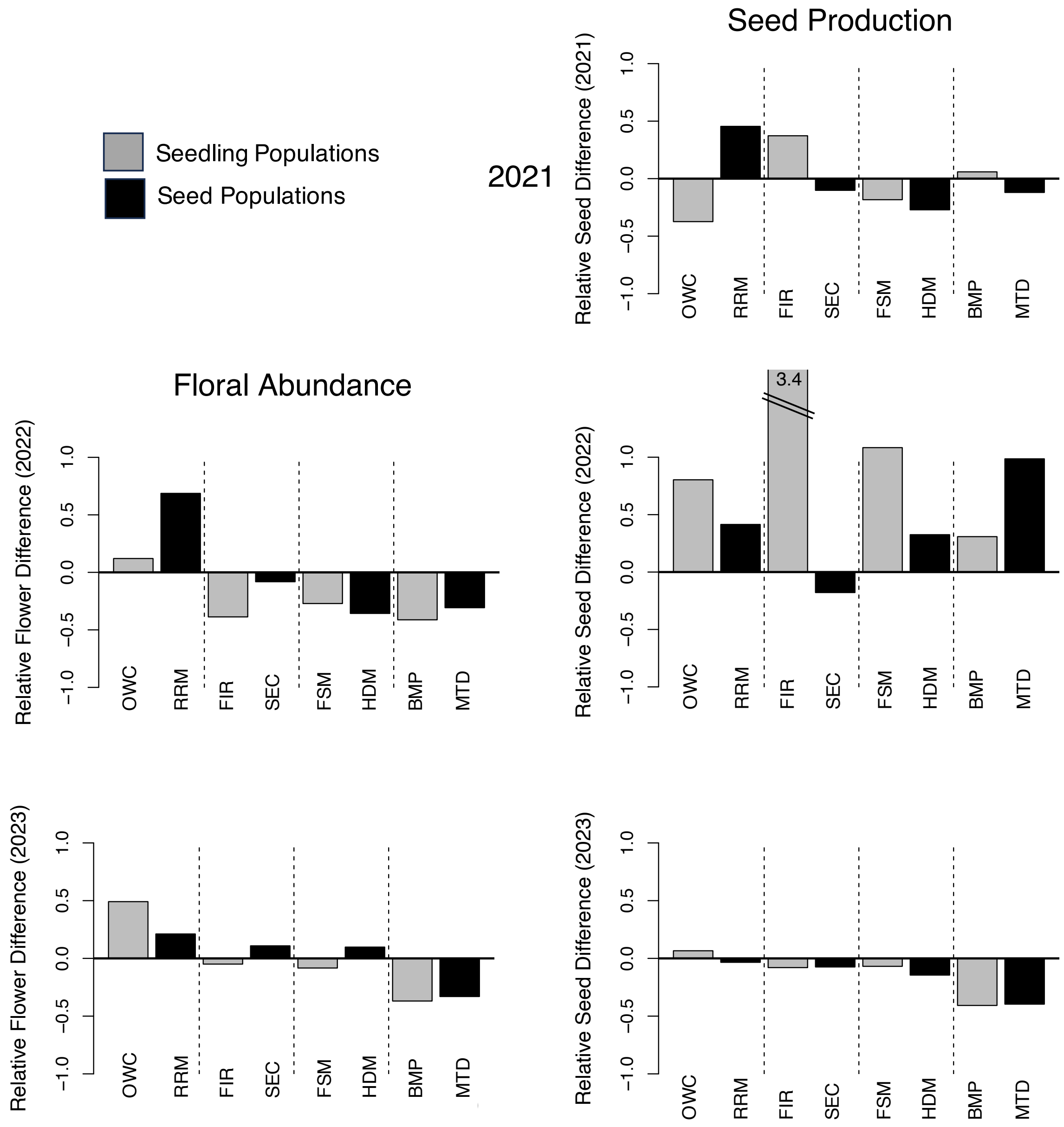


**Fig. S4 Relative changes in floral abundance and seed production across years following assisted gene flow**. Relative differences between control and treatment populations in each year were calculated: ($Treatment_{year}$ – $Treatment_{PreAGF}$) – ($Control_{year}$ – $Control_{PreAGF}$). $Treatment_{year}$ is the mean value in the focal population and focal year post-AGF, $Treatment_{PreAGF}$ is the mean value in focal population pre-AGF, $Control_{year}$ is the mean value in the control population and focal year post-AGF, and $Control_{AGF}$ is the mean value in control population pre-AGF. Bars above 0 indicate an increase in fitness relative to control populations post assisted gene flow. Grey bars represent populations where source seedlings were introduced while black bars represent populations where source seeds were introduced.

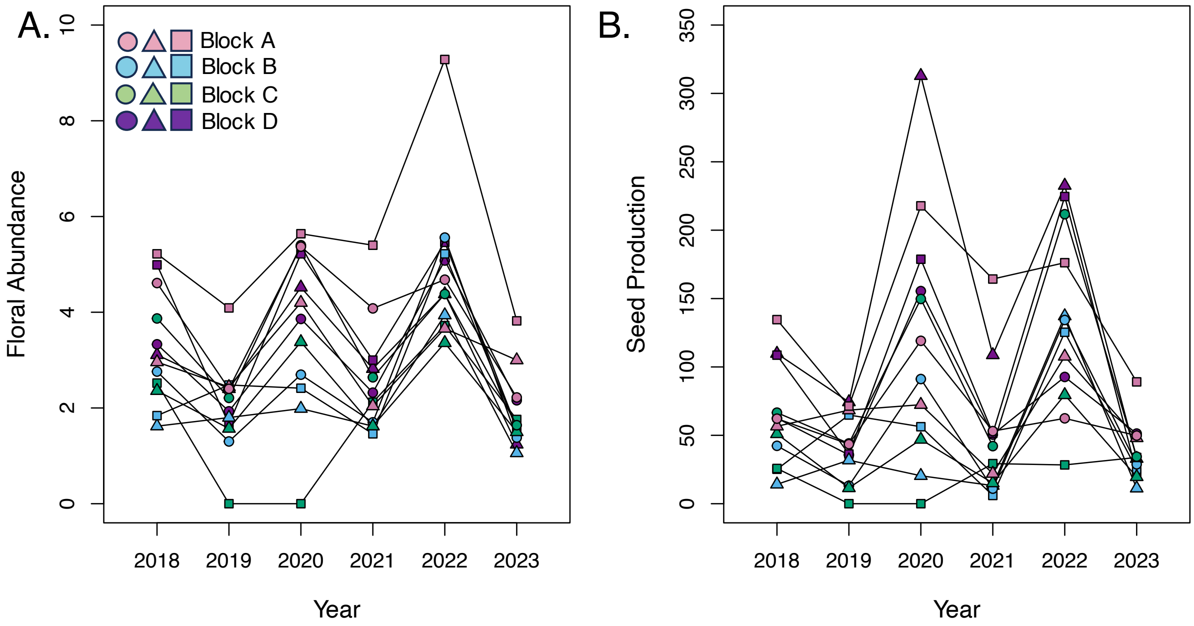


**Fig. S5. Temporal variation in fitness during the assisted gene flow experiment.** Variation in floral abundance (**A**) and seed production (**B**), respectively, across populations and years included in the experiment. Colors represent different experiment blocks and shapes represent treatment (local seeds = circles, source seeds = squares, source seedlings = triangles). Assisted gene flow occurred in 2021.

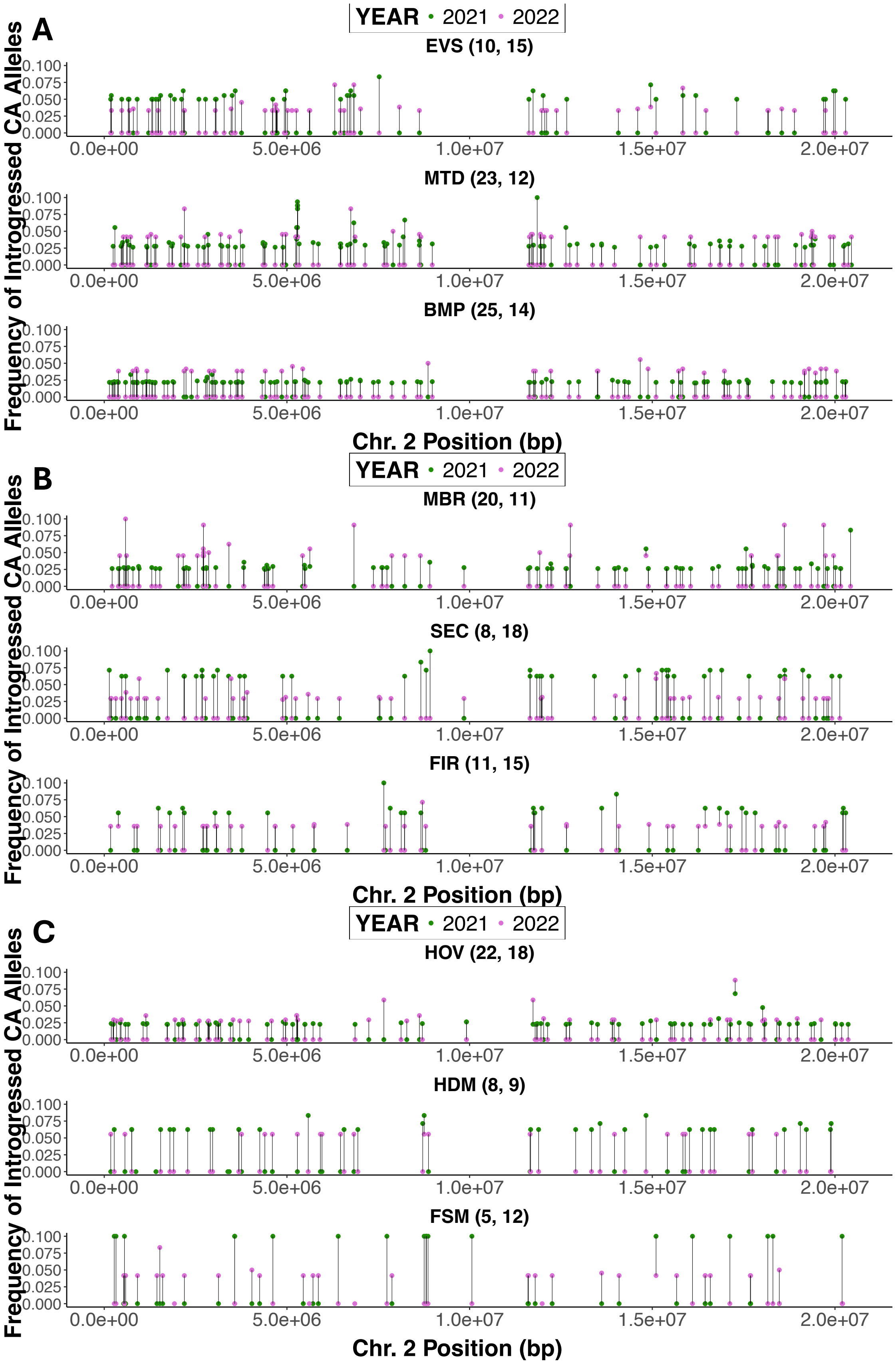
A
B
C
YEAR • 2021 • 2022
EVS (10, 15)
MTD (23, 12)
BMP (25, 14)
MBR (20, 11)
SEC (8, 18)
FIR (11, 15)
HOV (22, 18)
HDM (8, 9)
FSM (5, 12)
Frequency of Introgressed CA Alleles
Chr. 2 Position (bp)
0.000
0.025
0.050
0.075
0.100
0.0e+00
5.0e+06
1.0e+07
1.5e+07
2.0e+07

**Fig. S6 Limited introgression of California private alleles within target populations. A-C.** Allele frequencies of California private alleles across a representative chromosome (Chromosome 2) within Block D (**A**), Block B (**B**) and Block (**C**). Top graphs in each Block are from control populations (EVS, MBR, HOV), middle graphs are from populations receiving source seeds (MTD, SEC, HDM) and bottom graphs are from populations receiving source seedlings (BMP, FIR, FSM). Green and pink points represent the allele frequencies in 2021 and 2022, respectively. Within each subpanel, the top lollipop plot corresponds to the control population that received native seeds, the second to the population that received source seeds, and the bottom to the population that received source seedlings. Only loci that have evidence of introgression are represented for each population. Numbers in parentheses following population names reflect sample sizes for 2021 and 2022, respectively.

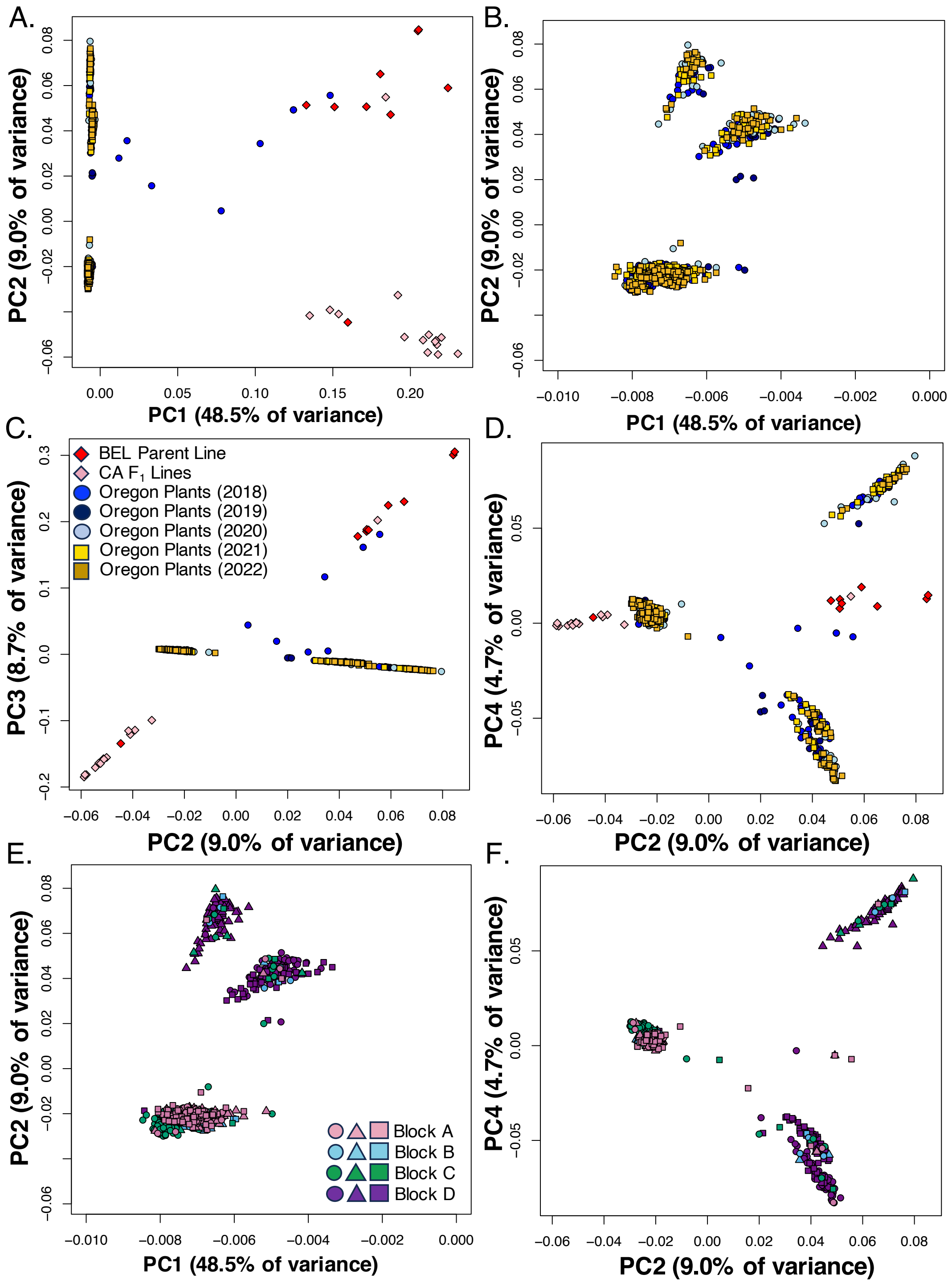


**Fig. S7 Populations structure across source and target monkeyflower populations. A.** Variation along PC1 and PC2 of all individuals from both source and target populations. A parent (BEL) and $F_1$ individuals from the source cross are represented by red and pink diamonds, respectively. All round points are Oregon target populations, with different colors representing different years. **B.** A zoomed-in scatter plot (note x-axis) of just the Oregon populations from panel A. **C-D.** Variation along PC3 and PC4 for source and target populations. **E.** Population structure among Oregon populations along PC1 and PC2 with colors and shapes representing different populations. **F.** Population structure among Oregon populations along PC2 and PC4.

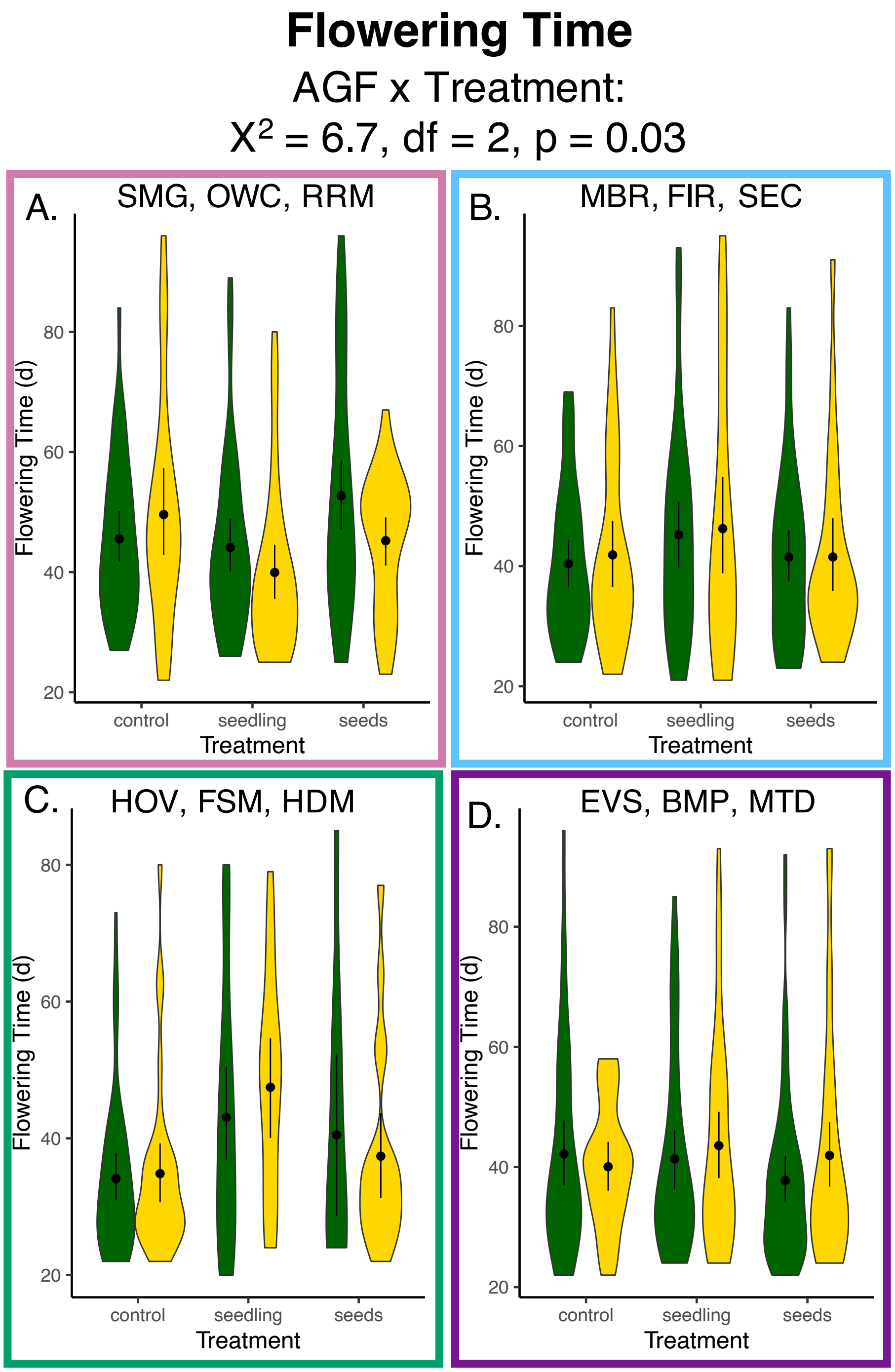


**Fig. S8 Variation between pre- and post-assisted gene flow in flowering time within a common garden by experimental block. A-D.** Violin plots depict variation in flowering time subset by experimental block from lowest to highest elevation block. Green plots represent the years before assisted gene flow, yellow plots represent the years following assisted gene flow. Points and whickers represent mean and standard error. Statistics below the main title correspond to a GLMM including all populations and years.

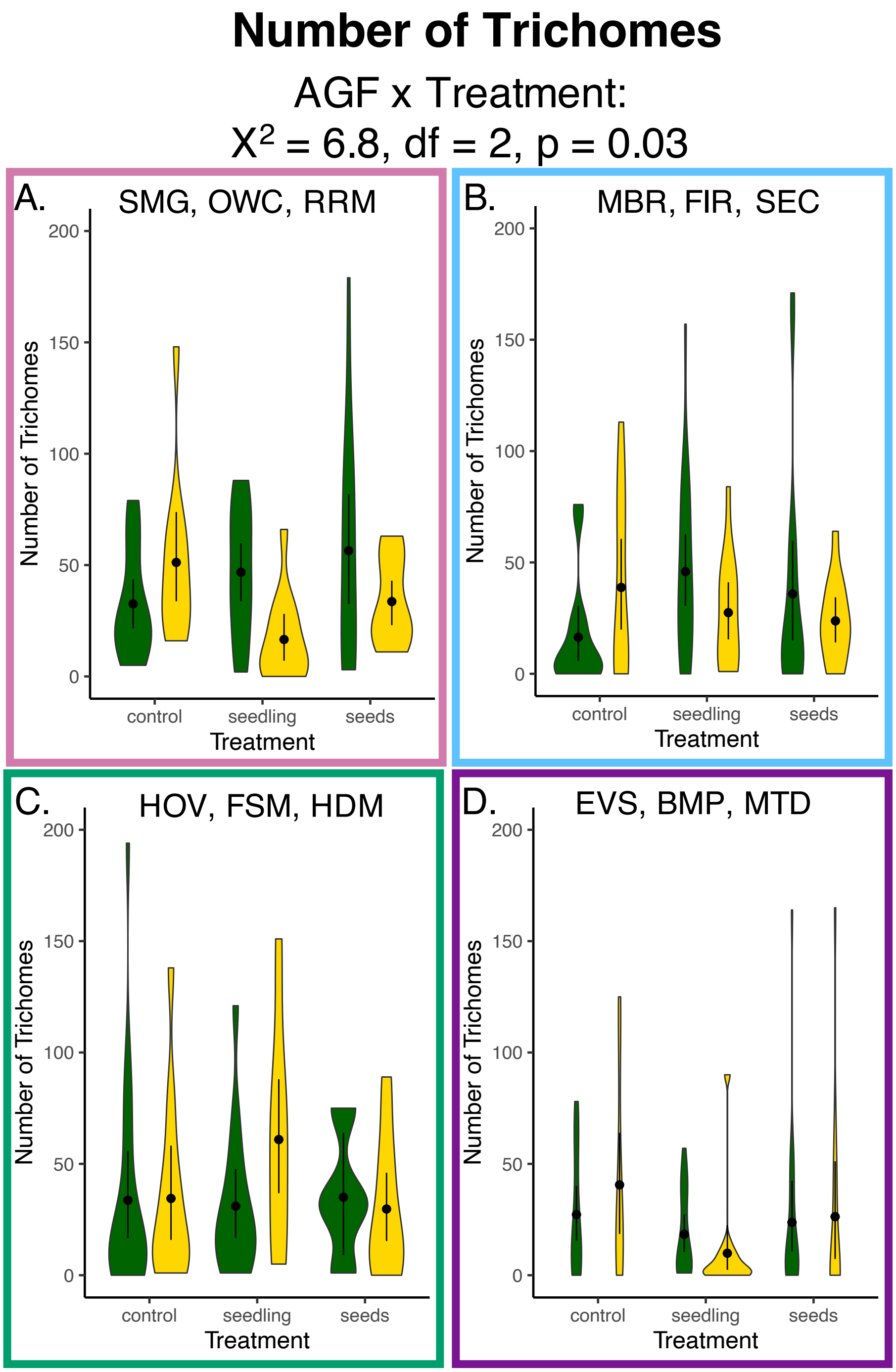


**Fig. S9 Variation between pre- and post-assisted gene flow in number of trichomes within a common garden by experimental block. A-D.** Violin plots depict variation in number of trichomes subset by experimental block from lowest to highest elevation block. Green plots represent the years before assisted gene flow, yellow plots represent the years following assisted gene flow. Points and whickers represent mean and standard error. Statistics below the main title correspond to the GLMM including all populations and years.
